# Oscillatory entrainment of the Frequency Following Response in auditory cortical and subcortical structures

**DOI:** 10.1101/2020.09.02.278119

**Authors:** Emily B.J. Coffey, Isabelle Arseneau-Bruneau, Xiaochen Zhang, Sylvain Baillet, Robert J. Zatorre

## Abstract

There is much debate about the existence and function of neural oscillatory entrainment mechanisms in the auditory system. The frequency-following response (FFR) is an index of neural periodicity encoding that can provide a vehicle to study entrainment in frequency ranges relevant to speech and music processing. Criteria for entrainment include the presence of post-stimulus oscillations and phase alignment between stimulus and endogenous activity. To test the hypothesis of entrainment, in experiment 1 we collected FFR data to a repeated syllable using magneto- (MEG) and electroencephalography in 20 healthy adults. We observed significant oscillatory activity after stimulus offset in auditory cortex and subcortical auditory nuclei, consistent with entrainment. In these structures the FFR fundamental frequency converged from a lower value over 100 ms to the stimulus frequency, consistent with phase alignment, and diverged to a lower value after offset, consistent with relaxation to a preferred frequency. In experiment 2, we tested how transitions between stimulus frequencies affected the MEG-FFR to a train of pure-tone pairs in 30 adults. We found that the FFR was affected by the frequency of the preceding tone for up to 40 ms at subcortical levels, and even longer durations at cortical levels. Our results suggest that oscillatory entrainment may be an integral part of periodic sound representation throughout the auditory neuraxis. The functional role of this mechanism is unknown, but it could serve as a fine-scale temporal predictor for frequency information, enhancing stability and reducing susceptibility to degradation that could be useful in real-life noisy environments.

## 1. Introduction

From brainstem to cortex, the auditory system responds in a synchronized manner to acoustical energy containing regular repeating elements. In low frequency ranges (<10 Hz), it may function to parse temporal structure of speech and music (Arnal and Giraud, 2012; Giraud and Poeppel, 2012; Nozaradan et al., 2011), integrate information (Schroeder et al., 2008), or enable selective attention (Schroeder and Lakatos, 2009; Golumbic et al., 2013). In higher frequency ranges associated with pitch information (80-400 Hz), frequency-following responses (FFRs) are observed (Kraus et al., 2017; Coffey et al., 2019; Skoe and Kraus, 2010; Krizman and Kraus, 2019).

Although rhythmic oscillations appear to be ubiquitous in the auditory system (Neymotin et al., 2020; Gourévitch et al., 2020), it is unclear whether such responses represent merely the summation of evoked responses to periodic sound elements, or arise from truly oscillatory intrinsic properties of the auditory system (Obleser and Kayser, 2019). The distinction is important to understanding their function (Gourévitch et al., 2020; Haegens and Golumbic, 2018), and fits into a bigger picture concerning the roles of intrinsic oscillations throughout the brain, in which they appear to play a role coordinating neuronal and network activity to support complex cognitive processes (Buzsáki and Draguhn, 2004; Thut et al., 2012).

The main aim of the present study is to clarify whether pitch encoding via the FFR involves a true entrainment mechanism. The FFR provides an excellent vehicle to address this question because its frequency content closely resembles that of the evoking stimulus, its strength correlates with behavioural and perceptual variables, and it is used widely in non-invasive human studies to quantify the quality of neural sound encoding (Kraus et al., 2017). Yet questions remain about its composition and origin (Bidelman, 2018; Tichko and Skoe, 2017; Coffey et al., 2016). According to the delay-based model, the FFR is generated via the summation of individual components from successive nuclei along the auditory neuraxis with increasing latencies (Tichko and Skoe, 2017; Gardi et al., 1979). In contrast, the oscillatory model suggests that the FFR is generated by neuronal circuitry with inherently oscillatory properties even at low levels (i.e. cochlea, inferior colliculus) (Lerud et al., 2019). The delay-based and oscillatory models make different predictions about auditory signal processing and perception.

In line with criteria for defining true neural entrainment (Haegens and Golumbic, 2018; Obleser and Kayser, 2019), we reasoned that if the auditory system’s representation of fine frequency information is oscillatory in nature, we should observe persistent activity at the stimulus frequency following stimulus offset (Notbohm et al., 2016). Furthermore, frequency representation should converge towards the stimulation frequency over time as information accrues through successive inputs to an oscillator (Thut et al., 2012; Giraud and Arnal, 2018).

Some fragmentary evidence in the literature exists to support the prediction of an FFR aftereffect following sound offset. Irvine (1986) measured single-unit recordings of the cat auditory nerve to periodic stimuli; Figure 8C of that work shows that instantaneous spike rates fluctuate for 3-4 cycles beyond the duration of the stimulus, at the stimulus frequency. More recently, Xu and Ye systematically recorded FFRs with EEG to stimuli with 6 durations which included 5-18 cycles of the fundamental frequency (Xu and Ye, 2015). Although the authors did not focus on the presence of an aftereffect, or on the relative durations of the responses, comparison of the number of cycles in the stimuli and the average responses (Figures 1 and 2 in Xu and Ye (2015)) suggests that there is indeed a post-stimulus response, which persists up to 3 cycles, dependent upon stimulus duration. A neural delay model cannot account for these observations. Regarding the second criterion of entrainment, to our knowledge, the prediction that the FFR frequency converges to the target frequency over time has not been tested.

**Figure 1:**
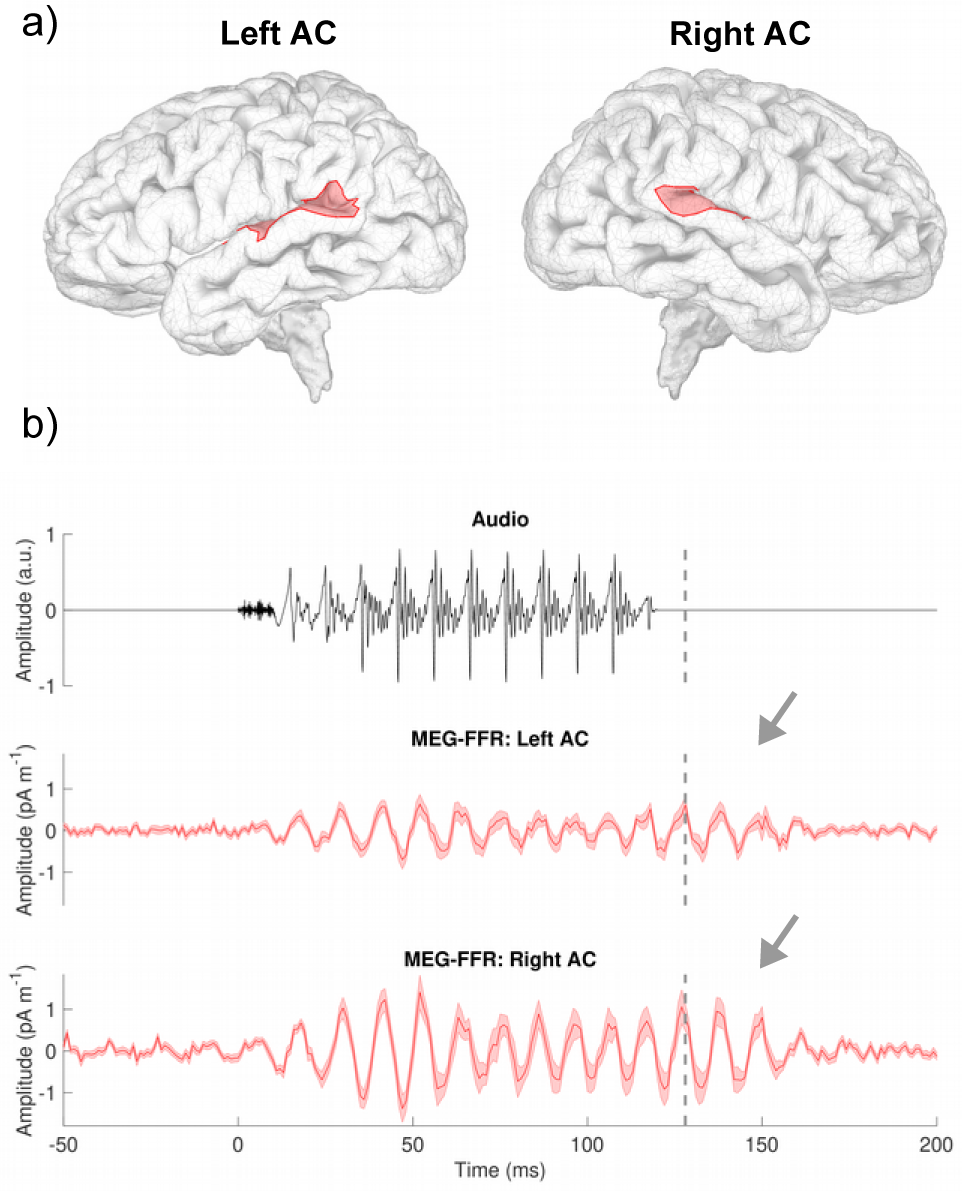
Activity in the left and right auditory cortex, regions illustrated in a sample subject in (a), continues for several cycles following stimulus offset (b, arrows). The grey dashed reference line indicates 10 ms following stimulus offset, which is believed to be the approximate feedforward transit time through the subcortical auditory system (Tichko and Skoe, 2017). Shaded bars represent standard error of the mean.

**Figure 2:**
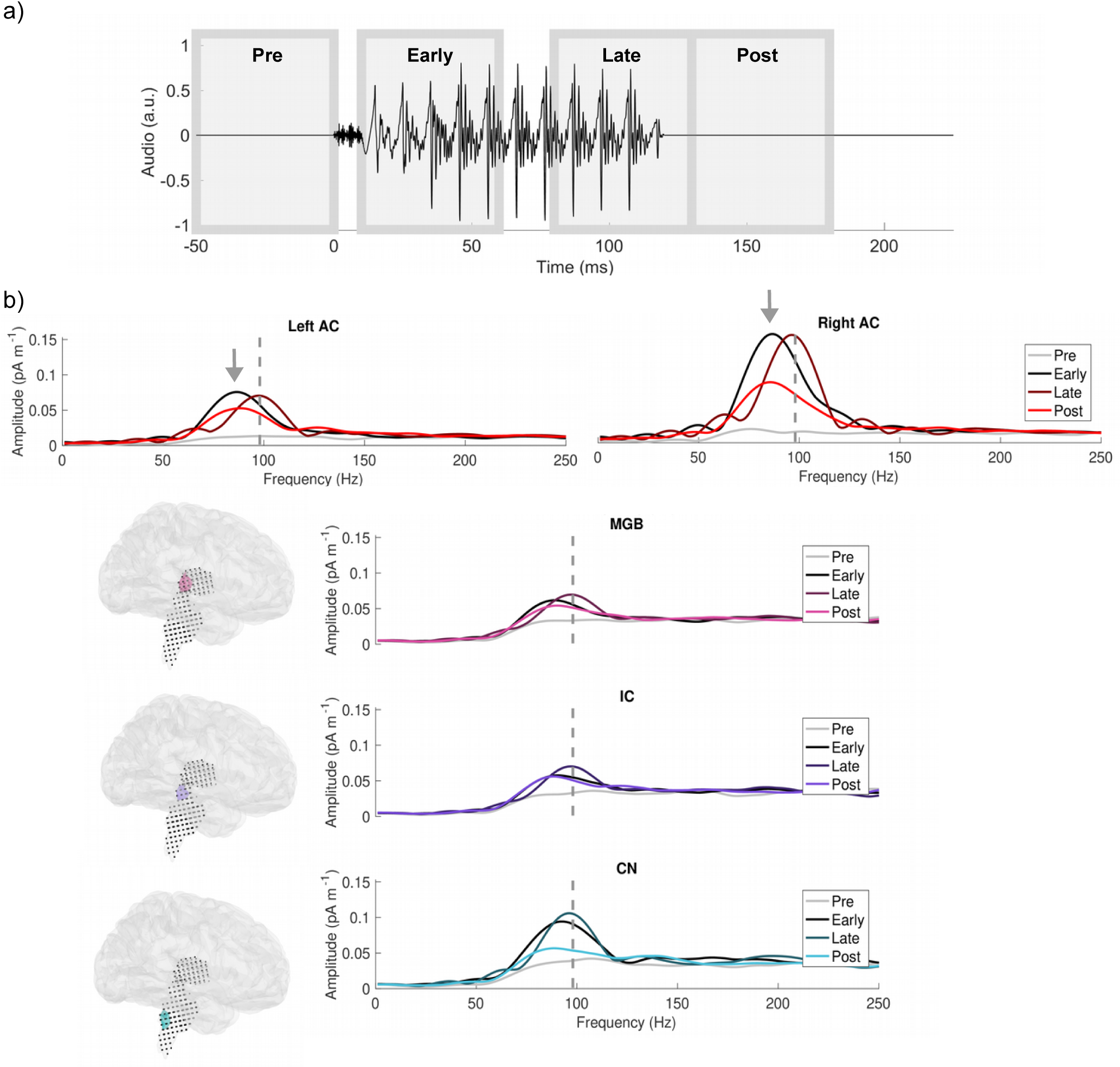
a) Schematic of the timing of the four 50 ms analysis windows relative to the auditory stimulation: prestimulation, during stimulation the first and second halves of the FFR, and post-stimulation. b) Mean amplitude spectra of each region of interest for each time period. The grey dashed line at 98 Hz denotes the fundamental frequency of the stimulus (F0). Arrows highlight the difference between peak frequency and stimulation frequency, which is most pronounced in the cortical regions, yet not during the late stimulation phase.

To evaluate evidence of oscillatory entrainment, we re-analyzed an existing dataset in which MEG/EEG-FFR was measured from healthy young adults listening to a speech syllable (Coffey et al., 2016), to test the following hypotheses (a) whether a post-stimulus aftereffect exists, (b) whether the frequency representation converges towards the stimulus frequency over time, and if so, (c) whether in the absence of continued stimulation it relaxes towards its original state (Obleser and Kayser, 2019). Finding positive results, we conducted a new MEG experiment to test whether a preceding stimulus affects the brain’s representation of an incoming stimulus. Our approach avoids the pitfalls of analysis which may produce artifactual oscillations (Gourévitch et al., 2020). Moreover, MEG allows for separation of cortical and subcortical sources of the FFR (Coffey et al., 2016) such that we can test whether these phenomena can be observed at all levels of the auditory system or not, which the existing literature does not address.

## 2. Experiment 1 Methods

### 2.1. Participants

Data from 20 neurologically healthy young adults included in a previous study (Coffey et al., 2016) were used in this study (mean age: 25.7 years; SD = 4.2; 12 female; all were right-handed and had normal or corrected-to-normal vision; ≤25 dB hearing level thresholds for frequencies between 500 and 4,000 Hz assessed by pure-tone audiometry; and no history of neurological disorders). Informed consent was obtained and all experimental procedures were approved by the Montreal Neurological Institute Research Ethics Board.

### 2.2. Stimulus presentation

The stimulus for the MEG/EEG recordings was a 120 ms synthesized speech syllable (/da/) with a fundamental frequency (F0) in the sustained vowel portion of 98 Hz. The stimulus was presented binaurally at 80 dB SPL, ∼14,000 times in alternating polarity, through Etymotic ER-3A insert earphones with foam tips (Etymotic Research). For five subjects, ∼11,000 epochs were collected due to time constraints. Stimulus onset asynchrony (SOA) was randomly selected between 195 and 205 ms from a normal distribution. To control for attention and reduce fidgeting, a silent wildlife documentary (Yellowstone: Battle for Life, BBC, 2009) was projected onto a screen at a comfortable distance from the subject’s face. This film was selected for being continuously visually appealing; subtitles were not provided in order to minimize saccades.

### 2.3. Neurophysiological recording and preprocessing

Two hundred and seventy-four channels of MEG (axial gradiometers), one channel of EEG data (Cz, 10–20 International System, averaged mastoid references), EOG and ECG, and one audio channel were simultaneously acquired using a CTF MEG System and its in-built EEG system (Omega 275, CTF Systems Inc.). All data were sampled at 12 kHz.

Data pre-processing was performed with Brainstorm (Tadel et al., 2011), and using custom Matlab scripts (The Mathworks Inc., MA, USA). Data were reanalyzed to extend the time window of analysis to 300 ms after stimulus offset, as the end of the response appears to have been truncated by the 150 ms window used previously in Coffey et al. (2016); however, analyses and figures will be limited to 200 ms which corresponds to the mean SOA. We took advantage of Brainstorm software developments in the interim to better standardize the location of regions of interest (ROIs), such that the centre of each region of interest was defined using standard space (MNI152) coordinates, transformed to the subjects’ T1-weighted MRI image, and visually inspected to confirm location. The coordinates of subcortical ROIs were: rCN: 8 −36 −38; lCN: −4 −36 −38; rIC: 6 −36 −10; lIC: −4 −36 −10; rMGB: 14 −30 6 and lMGB: −10 −32 6. Each ROI was between 0.4 and 0.5 cm^3^.

We used a distributed source modeling approach, in which the amplitudes of a large set of dipoles are used to map activity originating in multiple generator sites; these are constrained by spatial priors derived from each subject’s T1-weighted anatomical MRI scan (Gross et al., 2013; Baillet et al., 2001), from which cortical sources and subcortical structures were prepared using FreeSurfer (Fischl, 2012). Anatomical data were imported into Brainstorm, and the brainstem and thalamic structures were combined with the cortex surface to form the image support of MEG distributed sources: the mixed surface/volume model included a triangulation of the cortical surface (∼15,000 vertices), and brainstem and thalamus as a three-dimensional dipole grid (∼18,000 points). An overlapping-sphere head model was computed for each run; this forward model explains how an electric current flowing in the brain would be recorded at the level of the sensors, with fair accuracy (Tadel et al., 2011). A noise covariance matrix was computed from 1-min empty-room recordings taken before each session. The inverse imaging model estimates the distribution of brain currents that account for data recorded at the sensors. We computed the mininum-norm estimate (MNE) source distribution with unconstrained source orientations for each run using Brainstorm default parameters. The MNE source model is simple, robust to noise and model approximations, and very frequently used in literature (Hämäläinen, 2009). Source models for each run were averaged within subject.

The cortical ROIs were defined using the Destrieux atlas (Destrieux et al., 2010), by combining the regions labelled as ‘S temporal_transverse’, ‘G_temp_sup-Plantempo’, and ‘G_temp_sup-G_T_transv’ for the left and right hemispheres, respectively. This resulted in left and right auditory cortex regions (rAC, lAC) covering the posterior superior temporal gyrus.

We extracted a single timeseries of mean amplitude for each cortical ROI, and one for each of the three orientations (x,y,z) for the subcortical ROIs, which have unconstrained orientation. For analyses involving FFR-F0 strength and peak frequency, time windows of interest (described for specific analyses below) were obtained by first windowing the signal (5 ms raised cosine ramp), zero padding to 1 s to enable a 1 Hz frequency resolution, with subsequent fast Fourier transform, and rescaling by the proportion of signal length to zero padding. The spectra of the three orientations were then summed in the frequency domain to obtain the amplitude of each subject’s neurological response at the peak frequency close to the fundamental frequency, which was detected by an automatic script. Note that at the level of subcortical nuclei, magnetic fields emanating from activity of neural populations in the right and left sides are likely not spatially separable using the present techniques and nor do we address subcortical lateralization hypotheses herein; we therefore average them.

### 2.4. Data analysis

To evaluate the presence of a post-stimulus aftereffect, we first compared the strength of the oscillatory brain activity in a 20 Hz frequency band encompassing the fundamental frequency (88-108 Hz) in the post-offset period (130 to 180 ms) with the amplitude of the brain’s response at the same frequency during the pre-stimulus period (−50 to 0 ms), when no signal would be expected. For visual comparison, we calculated spectra averaged across subjects during the pre-stimulus window, two time periods of equivalent length (Early: 10 to 60 ms; Late: 80 to 130 ms) during stimulation, and during the post-stimulus window.

Non-parametric statistical tests are used throughout, as FFR amplitudes in a population tend to be non-normally distributed (Wilcoxon signed rank test; alpha = 0.05; one-sided). The post-offset period begins 10 ms post stimulus offset to account for approximate neural conduction delays to cortex (discussed in Tichko and Skoe (2017)). Values from subcortical structures were averaged across hemispheres, whereas data from the left and right auditory cortex were analyzed separately, for a total of 5 ROIs (lAC, rAC, MGB, IC, CN). Results were corrected for multiple comparisons using false discovery rate (FDR; (Benjamini et al., 2001)). Reasoning that the strength of the aftereffect should be somewhat related to the strength of the FFR, we used Spearman’s correlations (*r*_*s*_) to test for a positive relationship between the FFR amplitude in the Late window and the post-offset window across all levels (one-tailed, FDR-corrected). To investigate lateralization of the aftereffect at the cortical level, we compared the strength of the correlation between amplitude during vs. post stimulus on the right vs. left side, using Fisher’s exact test.

To investigate convergence of frequency accuracy, we first calculated the peak frequency of the averaged FFR responses by subject for each region of interest. To test whether the FFR peak frequency accuracy converges towards the stimulus’ frequency, we calculated the difference between the stimulus’ fundamental frequency (98 Hz) and each subject’s FFR in a 50 ms window at the beginning (10-60ms) and the end of (70-130ms) of the FFR (see Figure 2). Our primary interest was the MEG-FFR originating in the right auditory cortex, which has the highest amplitude and has been shown to correlate with individual differences in experience and perception (Coffey et al., 2016, 2017) and top-down processes such as attentional modulation (Hartmann and Weisz, 2019); however, we also conducted Wilcoxon signed-rank tests at each region of interest, correcting for multiple comparisons (FDR).

We then calculated peak frequencies over successive overlapping 50 ms windows (1 ms steps), and plotted mean and standard error of peak frequency for subjects whose SNR was greater than or equal to 5 (i.e. the amplitude of the fundamental was at least 5 times higher than the same frequency during the 50 ms silent baseline period preceding stimulus presentation), when at least 30 percent of subjects met the criterion. We plotted mean amplitude for subjects and datapoints reaching criteria, after first normalizing amplitude to that calculated during the baseline period over a ∼50 Hz frequency range centred on 98 Hz (73 to 122 Hz).

As a control, we confirmed that when subjected to the same analysis, the audio stimulus itself showed peaks precisely at 98 Hz in both the first and second half. Any difference in the FFR fundamental response found in these analyses are therefore due to signals generated in the brain and not inherent to the stimulus itself nor artifacts of the analysis.

When the stimulus ends, we expect an entrained oscillatory system with a preferred frequency that differs from that of the input to gradually relax towards that frequency. To test for oscillatory relaxation, we fit linear functions (least squares) to tracked frequencies during stimulation (80 to 120 ms post stimulus onset) and after stimulus offset (10 ms to 50 ms post stimulus offset) for each subject, and compared the resulting slopes for statistical differences at the group level using a Wilcoxon signed rank test (two-tailed). Such that all subjects had the same number of data points included in the linear fit, we did not exclude datapoints that did not meet an SNR criterion, as was applied in the previous analysis. Tests were conducted at each ROI, and reported p-values are corrected for multiple comparisons (FDR).

Because electroencephalography (EEG) is more commonly used to measure FFR (Coffey et al., 2019), and because MEG and EEG may differ in the degree to which they pick up different sources (Bidelman, 2018; Ross et al., 2020), we repeated the analyses described above in the simultaneously recorded single electrode (Cz referenced to mastoids) to investigate whether evidence of oscillatory phenomena might also be observed in EEG-FFR recordings.

## 3. Experiment 1 results

### 3.1. There is post-stimulation activity throughout the auditory system

In the MEG-FFRs from the left and right auditory cortex, the group averages clearly indicate the presence of continued activity following stimulus offset (see Figure 1). A conservative estimate of post-stimulus entrainment is used that excludes the first 10 ms following stimulus offset, during which time feed-forward transmission through the subcortical auditory system is expected (Tichko and Skoe, 2017). The timeseries is available for the cortex, which is modeled as a surface and yields a single timeseries per region of interest. We address the question of whether entrainment exists in subcortical regions, which are modeled as a volume and yield three timeseries per region, via spectral analysis.

At each level of the auditory system, there was significantly greater activity in the post-offset period at the stimulus’ fundamental frequency, as compared to activity at the same frequency in the pre-stimulus baseline period (lAC: *Z* = 3.70, *p* = .00027; rAC: *Z* = 3.73, *p* = .00047; MGB: *Z* = 3.32, *p* = .00074; IC: *Z* = 3.10, *p* = .0012; CN: *Z* = 2.80, *p* = .0026; p-values are FDR-adjusted).

The strength of the aftereffect was significantly correlated with the strength of the FFR in the Late window at the level of the rAC: *r*_*s*_ = 0.73, p = .00086 and IC: *r*_*s*_ = 0.49, *p* = .037; values trended in the same direction but did not reach significance at the level of the lAC: *r*_*s*_ = 0.34, *p* = .092; MGB: *r*_*s*_ = 0.32, *p* = 0.082; and CN: *r*_*s*_ = 0.42, *p* = .56. FDR-adjusted p-values are reported. The correlation between FFR and aftereffect strength was marginally stronger in the right as compared with left auditory cortex (*z* = 1.63, *p* = .051).

Figure 2 illustrates changes in spectral power pre-stimulus, during the FFR (early and late periods), and post-stimulus. The frequency of averaged peak amplitude differs from that of the stimulus during the early and post-stimulation periods, whereas during the late portion of the FFR, the FFR peak frequency matches that of the stimulus closely. This pattern is most pronounced at the cortical level.

### 3.2. Frequency accuracy increases with time

The difference between the stimulus’ fundamental frequency (98 Hz) and each subject’s FFR in a 50 ms window at the beginning (10 - 60ms) of the response was significantly greater than at the end of (70 - 130ms) of the response for all regions of interest except for the CN, which was marginal (see Table 1). These results suggest that frequency tracking improves over time, converging towards the stimulation frequency as information is accrued.

**Table 1:**
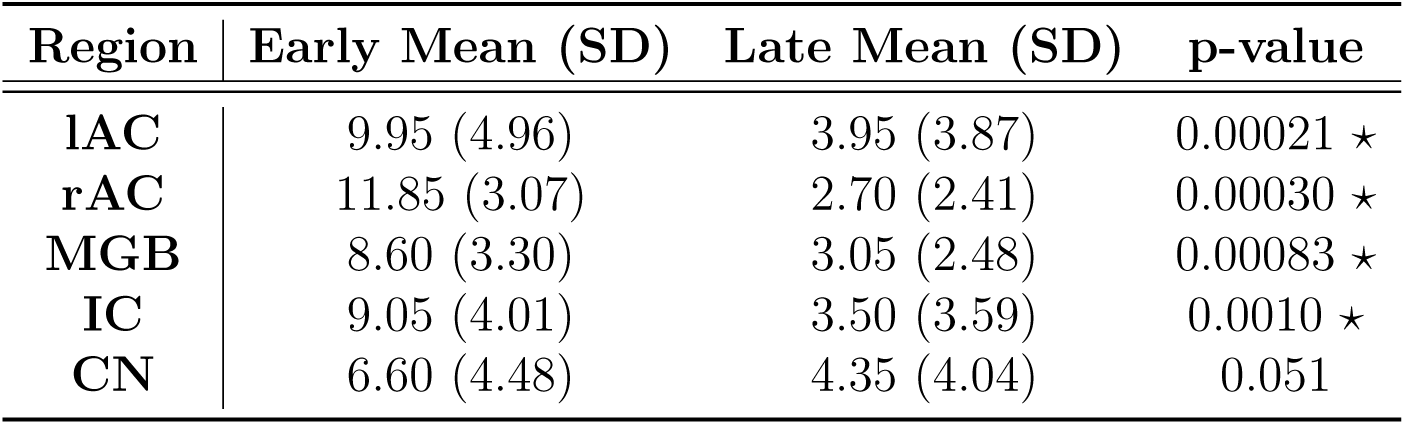
Difference between peak frequency and the 98 Hz stimulation frequency, during the early (10 - 60 ms) and late (80 - 130 ms) portions of the FFR. The mean and standard deviation (SD) are in Hz (absolute value of difference from F0). Statistically significant comparisons are indicated with an asterisk (⋆), and are FDR corrected.

A finer-grained analysis of peak-frequency in 50 ms overlapping windows reveals a pattern of convergence towards the fundamental frequency of the stimulus, in both the left and right auditory cortex (Figure 3). Mean peak frequencies are plotted for each window, for subjects who showed a clear peak close to the stimulus F0 (as defined as 5 times higher than the amplitude of the same frequency found in the pre-stimulus period). The FFR reaches the 98 Hz stimulation frequency about 100 ms following stimulation onset (∼9 cycles of the vowel’s F0). After sound offset, the mean peak frequency decreases. Tracking results are unstable at the beginning and end of the analysis window, indicated by higher tracking variability, when FFR amplitude is low and few subjects showed clear spectral peaks.

**Figure 3:**
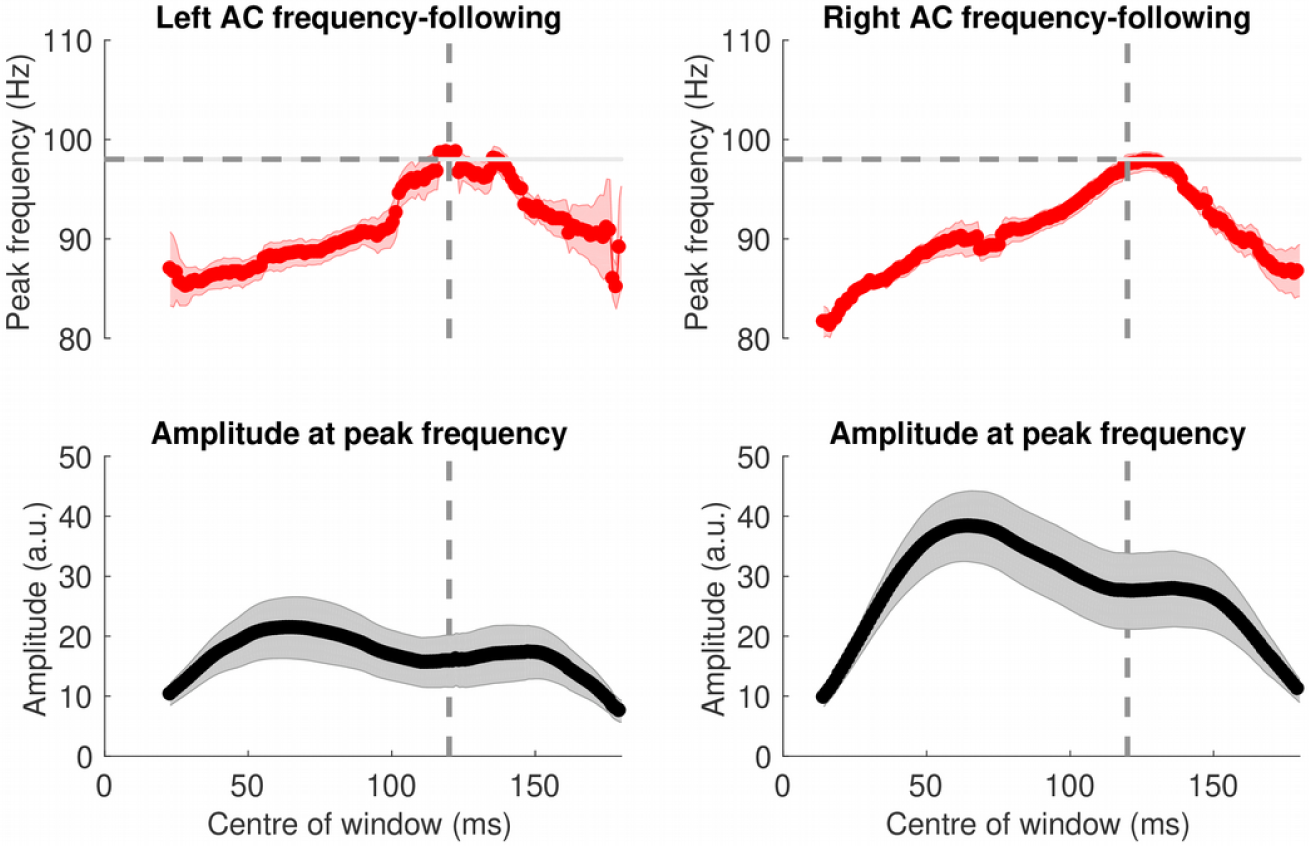
Frequency tracking in the left and right auditory cortex (top), and the corresponding normalized amplitude of the detected peaks (bottom), over time. Data are plotted for 50 ms windows in which at least 30 percent of subjects had peaks that reached a +5 SNR threshold with respect to baseline. Centre of window is reported relative to stimulus onset. The horizontal and vertical dashed lines represent stimulus F0 and offset, respectively. Shaded bars represent standard error.

In the subcortical structures, the frequency tracking accuracy also increases towards the end of the stimulus, as it does for the cortex, but as the initial frequency is closer to the stimulus F0, a less pronounced pattern of convergence is observed (see Figure 4).

**Figure 4:**
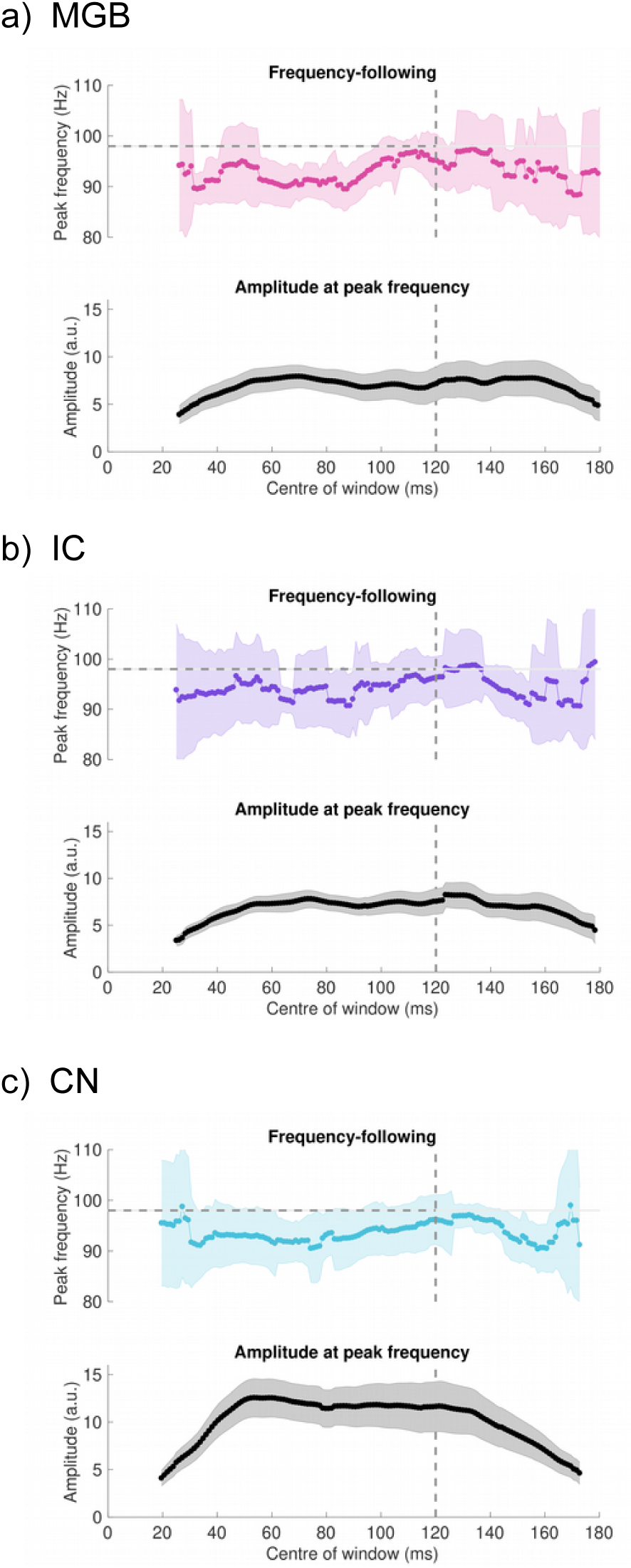
Frequency tracking and normalized amplitude of detected peaks in the a) medial geniculate nucleus (MGB), b) inferior colliculus (IC) and c) cochlear nucleus (CN). Data are plotted for 50 ms windows in which at least 30 percent of subjects had peaks that reached a +2 SNR threshold with respect to baseline. The horizontal and vertical dashed lines represent stimulus F0 and offset, respectively. Shaded bars represent standard error.

### 3.3. Tracked frequency relaxes after external input ends

Positive relationships between time and tracked frequency before stimulus offset and the negative relationship following offset indicates first convergence towards 98 Hz, followed by a return to lower frequencies. At the rAC, the mean slope of tracked frequency during stimulus presentation was 0.20 (*SD* : 0.14), and after stimulus offset was −0.31 (*SD* : 0.18). The pattern was significant at the rAC level (*Z* = 3.91, *p* = .00044). A similar pattern of results was found at the lAC (mean during: 0.17, *SD* : 0.24; mean afterwards: −0.21, *SD* : 0.30; *Z* = 3.29, *p* = .0025). At sub-cortical levels, slope pre- vs. post-stimulus offset was significantly different at the MGB (mean during: 0.21, *SD* : 0.31; mean afterwards: −0.14, *SD* : 0.38; *Z* = 2.31, *p* = .034), IC (mean during: 0.15, *SD* : 0.29; mean afterwards: −0.15, *SD* : 0.41; *Z* = 2.12, *p* = .033), and CN (mean during: 0.14, *SD* : 0.22; mean afterwards: −0.11, *SD* : 0.35; *Z* = 2.20, *p* = .035), FDR adjusted p-values are reported.

### 3.4. Evidence for oscillatory phenomena in the EEG-FFR

In the single-channel EEG recording, as observed in MEG, there was significantly greater activity in the post-offset period close to the stimulus’ fundamental frequency, as compared to activity at the same frequency band in the pre-stimulus baseline period (*Z* = 3.92, *p* = .00014; see Figure 5a).

**Figure 5:**
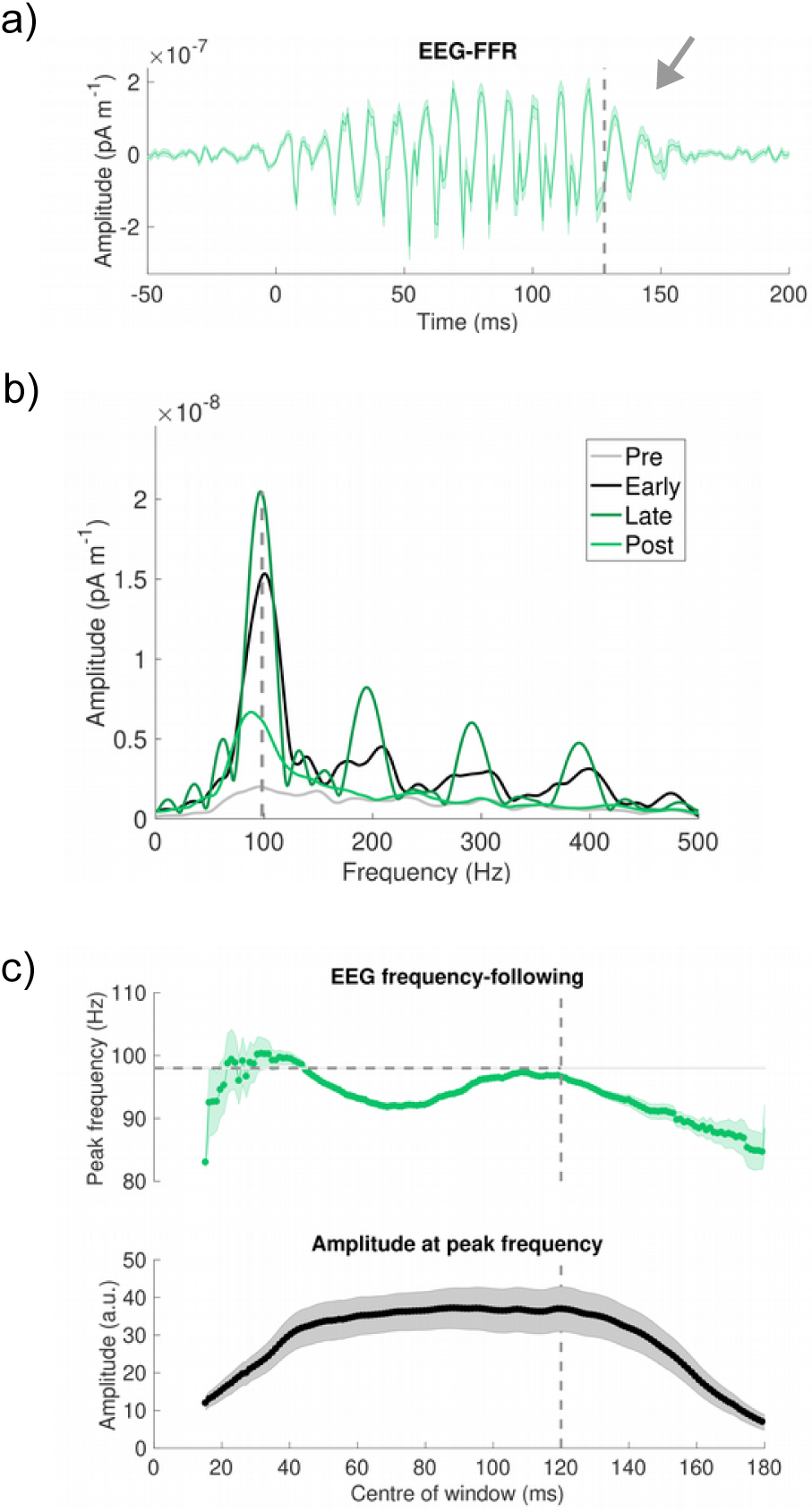
Evidence for oscillatory phenomena in EEG. a) Activity in the EEG recording continues for several cycles after stimulus offset (a, arrow). b) Mean amplitude spectra for the EEG-FFR for each time period (as in Figure 2a). In contrast to the AC, EEG-FFR does not exhibit a significant difference in frequency representation in early vs. late FFR periods. c) Frequency tracking in EEG trace (top), and the corresponding normalized amplitude of the detected peaks (bottom), over time. Data are plotted for 50 ms windows in which at least 30 percent of subjects had peaks that reached a +5 SNR threshold with respect to baseline. The horizontal and vertical dashed lines represent stimulus F0 and offset, respectively. Shaded bars represent standard error.

In contrast to the MEG data from regions of interests, the accuracy of frequency representation was not significantly better during the late FFR period (mean difference from stimulus F0: 2.95, *SD* : 3.02) than the early FFR period (mean: 2.10, *SD* : 1.59; *Z* = 1.36, *p* = 0.087); nor did the averaged spectral peak differ between the early and late period (see Figure 5b). The frequency tracking analysis (Figure 5c) suggests that following the stimulus offset, the frequency trends towards lower frequencies, as observed in the AC (but less so in the subcortical MEG-FFRs). Qualitatively, the tracking results of the EEG signal do not resemble those of any single one of MEG ROI signals, further supporting the proposal that the EEG-FFR is a composite signal that may represent different mixtures of sources over its duration (Coffey et al., 2019; Tichko and Skoe, 2017).

As noted in previous work (Coffey et al., 2016), harmonic components (i.e. 196, 294, and 392 Hz) are clearer in the EEG than MEG signal. Interestingly, the harmonics appear to be strongly represented in the late FFR period but are do not appear in the post-FFR period, suggesting that the oscillatory phenomenon may be related only to representation of the stimulus’ envelope rather than its high frequency fine structure. Although these results raise further questions about the interactions between structures, and the representation of subcortical and cortical generators in the compound EEG-FFR signal recorded at the scalp, our results generally support the involvement of oscillatory dynamics in the brain’s encoding and representation of periodic acoustical information.

## 4. Experiment 2

Finding positive evidence for oscillatory phenomena, we designed a new study to test the prediction that the frequency representation of an incoming stimulus would be influenced by the frequency content of an immediately preceding stimulus. We created a stimulus sequence in which three short complex tones with different fundamental frequencies (tones A: 98 Hz, B: 131 Hz, and C: 147 Hz) were presented in a continuous stream (with equal transitional probabilities). This arrangement allowed us to examine how the recent exposure to the higher-pitched tones B and C affected the representation of tone A, which has the same pitch as the stimulus in Experiment 1 and is robustly represented at all system levels.

In Experiment 1, we had observed an unexpected phenomenon: that convergence at the cortical level occurred consistently from lower rather than random frequencies (i.e. <98 Hz; Figure 3), when sound was presented following a short period of silence. This observation is consistent with the existence of a preferred or resonant frequency that is somewhat lower than the 98 Hz fundamental, but can be progressively entrained to 98 Hz over about 100 ms (∼9 cycles for tone A). In Experiment 2, we hypothesized that after 9 cycles of entraining to a higher-pitched tone, convergence to Tone A would occur from higher, rather than lower frequencies as observed in Experiment 1, and that differences in sensory history would affect the representation of a stimulus.

## 5. Experiment 2 Methods

### 5.1. Participants

Thirty neurologically healthy young adults (mean age: 25.8; SD = 4.8; 15 female; all were right-handed and had <25 dB hearing level thresholds for frequencies between 500 and 4,000 Hz assessed by pure-tone audiometry; and no history of neurological disorders).

### 5.2. Experimental procedure

Subjects were recruited to participate in a three-session experiment in which the effect of transcranial magnetic stimulation (TMS) to the right auditory cortex 5 mins prior to FFR recording was assessed. Here we report only the MEG data from the sham TMS condition, in which the TMS device was discharged perpendicular to the head as a control condition for the other sessions (i.e. no magnetic stimulation of the brain, but the subjects were aware of clicks). Written informed consent was obtained and all experimental procedures were approved by the Montreal Neurological Institute Research Ethics Board.

### 5.3. Stimulus presentation

Each run included a random sequence of 3 tones which were comprised of a sine wave at the fundamental frequency, added to sine waves at the frequencies of its 2^nd^ to 4^th^ harmonics (see Figure 6). Each tone includes 9 cycles of its fundamental frequency, such that the sine waves of each harmonic begin and end at a zero phase; therefore the stimuli are of different durations. The fundamentals of the tones were as follows (corresponding Western musical scale value given in parentheses): tone A (G2): 98 Hz (period is 10.2ms, duration 91.83 ms); tone B (C3): 131 Hz (period is 7.7ms, duration 68.83 ms); and tone C (D3): 147 Hz (period is 6.8 ms, duration 61.17 ms). A silent gap the same length as tone C was occasionally presented between two tone C presentations for an unrelated research question. A 5 ms raised cosine ramp was applied to each stimulus to avoid clicks.

**Figure 6:**
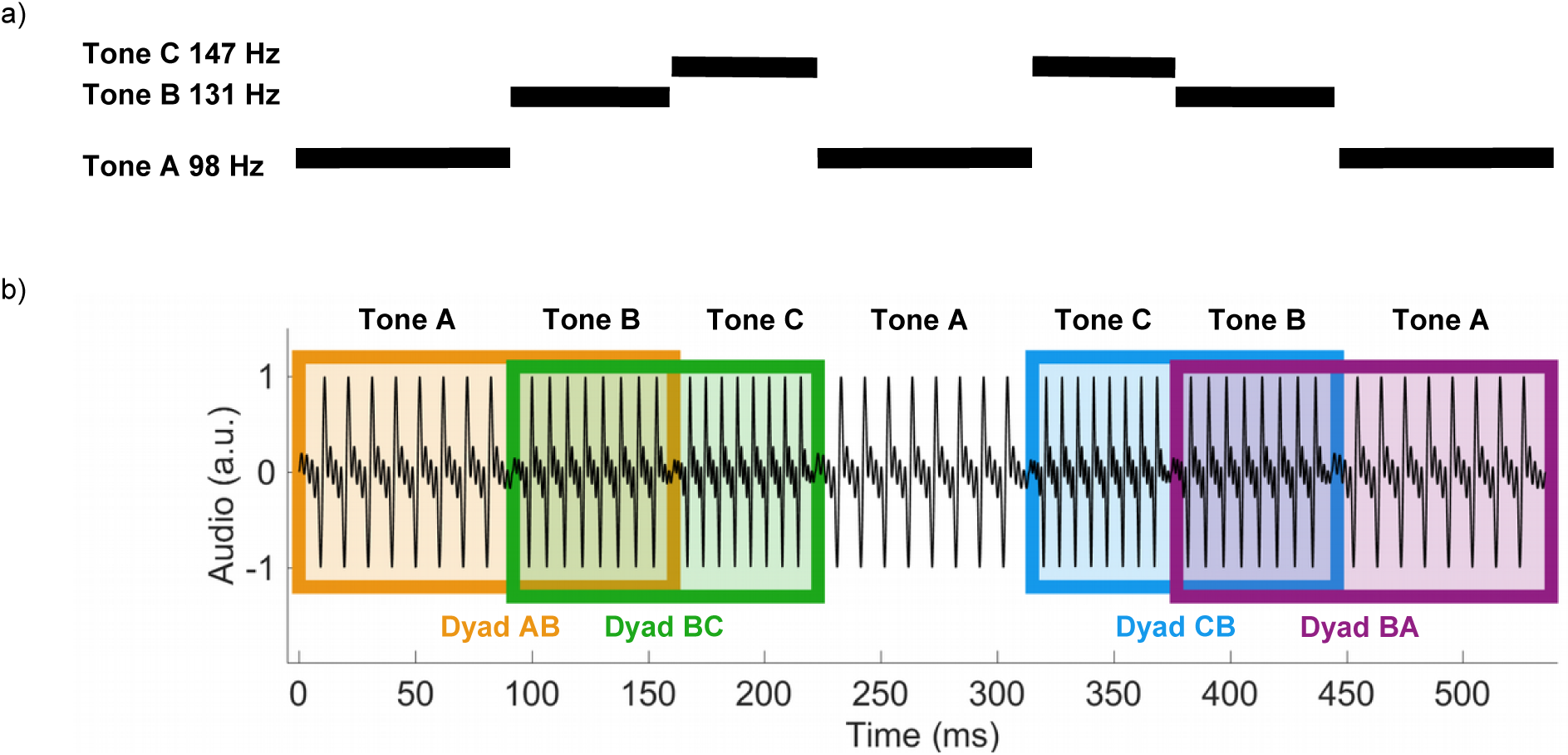
Audio stimulation used in Experiment 2, in which three tones are presented in a continuous stream. a) The relative pitch of the three tones, shown schematically in a representative 500 ms segment. The duration of each tone is equivalent to 9 cycles of its fundamental frequency, and includes the fundamental frequency and 2^nd^-4^th^. b) Tone pair dyads are defined within the continuous stream for analysis, such that the relative effects of higher or lower-pitch preceding tones may be studied.

### 5.4. Neurophysiological recording and preprocessing

MEG recording was identical to that described in Experiment 1, except that data were sampled at 6 kHz, which is sufficient to capture the fundamental frequencies which are the focus of these analyses. Pre-processing was also as described above, except that data were bandpass-filtered (80 - 450 Hz), downsampled to 1000 Hz, and epoched in windows of −50 ms to 351 ms relative the onset of the first of a pair of transition dyads (e.g. AB, BC), and later trimmed to to the portion of data relevant to each analysis, as will be further described in each section. In total, each dyad occurred slightly more than 2000 times. Whole-cap EEG was simultaneously measured with an external system (BrainAmp MR, Brain Products GmbH, Gilching, Germany) to address research questions not reported here.

### 5.5. Data analysis

All three tones have fixed pitches. If all three tones are represented with high fidelity in the brain, we would expect discrete peaks in the spectra at the fundamental frequencies of those tones (i.e., at 98, 131, and 147 Hz), to the extent that levels of the auditory hierarchy are capable of robustly representing those frequencies within the stimulation time-frame. If changing between frequency representations occurs gradually over multiple cycles as suggested by Experiment 1, we expect broad peaks as the representations blur together. To observe the frequency representation across regions of interest, we first calculated mean spectra across responses to all stimuli at each level of the auditory system using a Fourier transform, averaged across subjects for each region of interest.

In Experiment 1, the right auditory cortex showed a stronger, clearer FFR than the left. To test whether the previously observed result was replicable, we evaluated if the amplitude of the right AC for the tone A was statistically greater than that of the left (Wilcoxon signed rank test, one sided).

We tracked the frequency representation (as described in Experiment 1, using a 50 ms sliding window) during the presentation of tone A that occurred after tones B and C. Because each tone does not have a preceding silent period that can be used as a baseline for an SNR threshold, we instead calculated mean amplitude within the 250-450 Hz frequency range for each subject during tone presentation, as the FFR spectra have a relatively flat frequency response within this range. Data points were included in the group average that exceeded the high frequency range mean by a factor for 2. Data were smoothed with a 5 point moving average filter for visualization purposes.

To fairly test whether frequency representation convergences from higher frequencies to tone A’s fundamental, we must avoid including tracking data points that are derived from windows that overlap with the preceding stimulus. Therefore we used a short tracking window (30 ms) to allow for multiple sliding windows, and included window centres starting at 25 ms (i.e. window half width of 15 ms plus the 10 ms period following the transition to avoid the possibility of contaminating the response from neural activity of the preceding stimulus). As in the slope-based analysis in Experiment 1, we do not exclude data points on the basis of SNR, such that all subjects and brain regions have the same number of points. We then fit linear functions to the tracked frequencies of 25 to 85 ms, and tested whether slopes are less than zero, indicating convergence from higher to lower frequencies, using a Wilcoxon signed rank test (FDR-corrected for multiple comparisons across ROIs, within each condition).

To observe the effect of preceding stimuli on phase and amplitude in the time domain, we bandpass filtered (80-150 Hz) the timeseries for dyads AB, CB, BA, and BC, and calculated a group average and standard error aligned to the transitions between the tones for each pair, for qualitative visual inspection. Because the expected aftereffect lasts 3-4 cycles, a 10 ms buffer period for neural delay should be maintained, and a window of at least a few cycles is needed to accurately quantify oscillatory activity in a spectral analysis, it is difficult to cleanly quantify the effects of the aftereffect by comparing potions of dyad pairs with different transitions in the frequency domain. We conducted post-hoc analyses on amplitude of Tone B, focusing on the AB vs. CB comparison, testing the hypothesis that the amplitude of B would be higher when preceded by A as opposed to C. We did not test the effect of tone B on tones A and C because in that case the prediction is that A and C are similar to one another, hence essentially the null hypothesis which cannot be tested. We defined two discrete 30 ms time windows that capture information about the brain’s response to sound starting 10 and 40 ms after the transition to B (‘early’ and ‘late’) for further analysis, illustrated in Figure 8. Instead of using the amplitude of spectral peaks, which can be less reliably detected in such short time windows particularly for deeper regions, we averaged the amplitude values within a frequency band covering all tone fundamental frequencies (i.e. 80 - 160 Hz) and computed the relative difference between tone B’s amplitude when it occurred following tone A vs. tone C for each subject. This measure captures the relative difference in amplitude of frequency representation across conditions, even if the brain’s response has not fully converged to the stimulus’s fundamental frequency.

## 6. Experiment 2 Results

### 6.1. Replication of right-lateralization of cortical FFR

We confirmed that the amplitude of the right AC for the Tone A (mean: 0.060, SD: 0.031) was statistically greater than that of the left (mean: 0.037, SD: 0.016); Z = 4.1, P <0.0001), replicating the finding of strong lateralization of amplitude in the cortical MEG-FFR, in this larger sample. We hereafter focus on the right auditory cortex response based on the observation from Experiment 1 that patterns of convergence and the duration of the aftereffect in left and right auditory cortex was similar (refer to Figure 1).

### 6.2. Frequency tracking is modulated by preceding frequencies

Results of frequency tracking in the right auditory cortex for tone A following either C or B, which are both higher-frequency, are presented in Figure 7.

**Figure 7:**
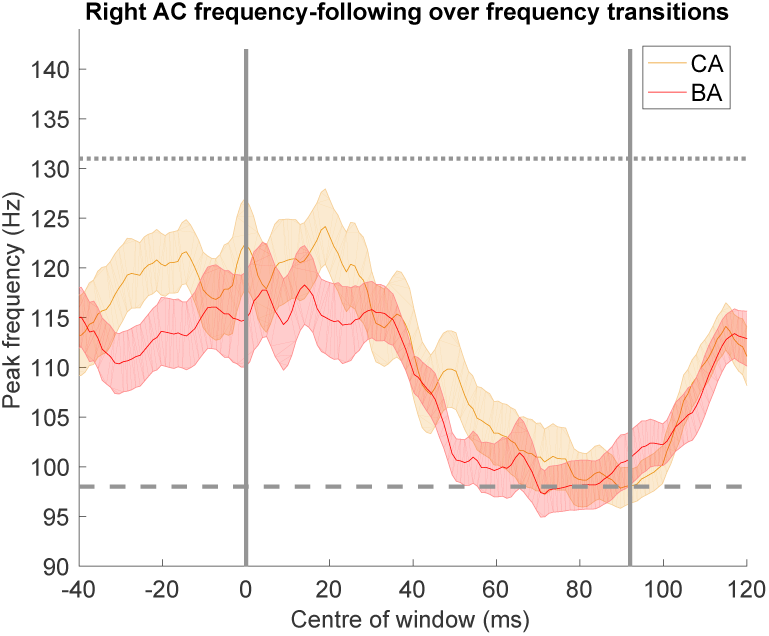
Frequency tracking in the right auditory cortex for tone dyads BA and CA, which both start with a higher-frequency tone. Although tracking results at the end of C and B imply an incomplete convergence to their stimulation frequencies by tone offset (62 and 69 ms, respectively), the pattern of results after transition to tone A (98 Hz, dashed horizontal line) demonstrates frequency tracking convergence from higher frequencies. Data are plotted for 50 ms windows for subjects who had peaks that reached a +2 SNR threshold in each window (relative to mean amplitude averaged in a 250 - 450 Hz frequency band). Time is reported relative to onset of tone A, and vertical lines mark tone transitions to and from tone A. Shaded bars represent standard error.

We evaluated the hypothesis that tone A representation was affected by having been preceded by tone B, by testing whether the slope of frequency tracking was less than zero, indicating convergence from higher frequencies to tone A’s 98 Hz fundamental. At the rAC, the mean slope of tracked frequency during tone A stimulus presentation was −0.21 (SD: 0.36), which was significantly below zero (Z = −3.24, *p* = .0030) as were results at the lAC (mean slope: −0.30, SD: 0.36; Z = −3.67, *p* = .00030). At subcortical levels, the signals of which have lower SNR, the success of tracking results appeared unstable across participants, suggesting that the design and data did not allow for reliable estimate of frequency needed for this analysis. Results are reported for completeness: MGB mean slope: 0.12, SD: 0.37; Z = 1.49, *p* = .43), IC (mean slope: −0.12, SD: 0.39; Z = −1.49, *p* = .085), and at the CN (mean slope: 0.14, SD: 0.35; Z = −2.05, *p* = .034).

We repeated the process for tone A when followed by tone C, finding a similar pattern of results at the cortical level; at the rAC, the mean slope of tracked frequency during tone A stimulus presentation was −0.31 (SD: 0.44; Z = −3.36, *p* = .0019); and at the lAC (mean slope: −0.22, SD: 0.41; Z = −2.66, *p* = .0098). Subcortical results are reported for completeness: MGB (mean slope: 0.06, SD: 0.38; Z = 0.46, *p* = .22), IC (mean: 0.06 slope: 0.38, SD: 0.xx; Z = 0.46, *p* = .28), and at the CN (mean slope: −0.037, SD: 0.41; Z = −0.79, *p* = .21). Reported P-values are FDR-adjusted within each analysis.

### 6.3. Effect of preceding events on sound representation

To further investigate if and where convergence towards stimulation frequency occurs progressively in a process of oscillatory entrainment, we examined the mean timeseries of the transition point between converging (AB and CB), and diverging dyads (BA vs BC). Qualitatively, it appears that after the AB and CB transitions, the phase converges somewhat, whereas in the BA vs BC comparison, the signal appears similar post transition suggesting a lingering effect of B. However, although phase differences are suggestive in the grand average, they appear highly variable at the subject level. The grand averages suggest that persistent differences may be more clearly manifested in measures of amplitude; namely, the amplitude of the FFR to tone B appears higher after the higher-amplitude A FFR than after the lower-ampliutde C FFR (Figure 8, top). Conversely, the brain’s response to tone A and C and appear highly similar when both preceded by Tone B, despite that the stimuli differ in frequency by 49 Hz (Figure 8, bottom).

**Figure 8:**
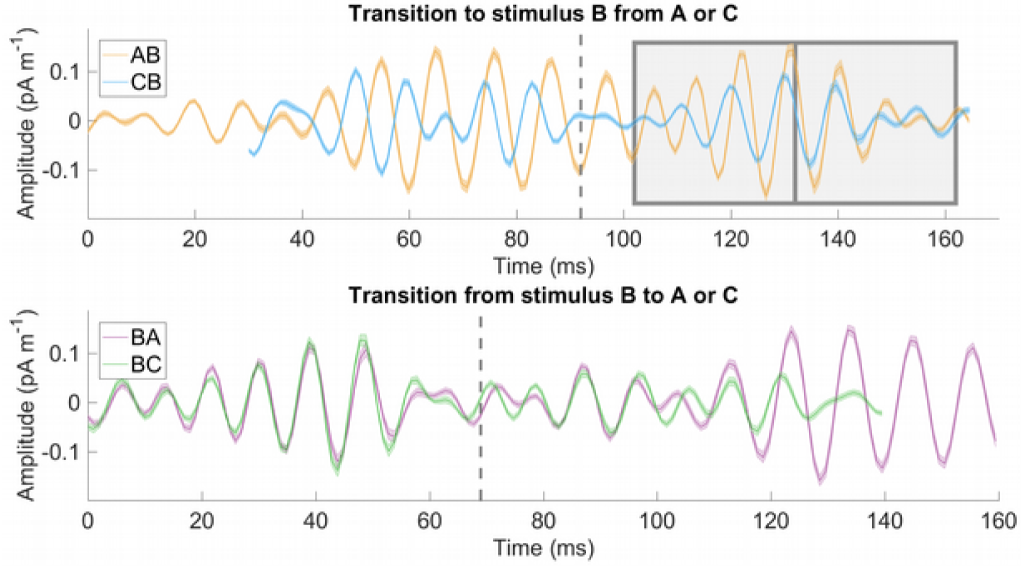
Activity in the right auditory cortex in response to pairs of tones, aligned at the transition point between tone dyads (dashed lines). In the transition to the same stimulus (top), the cortical response to tone B appears to differ, whereas after transition from tone B (bottom), both traces resemble one another despite a 50 Hz difference in stimulus frequency. Early (10 - 40 ms) and late (40 - 70 ms) windows are marked on the first dyad for further analysis. Shaded areas represent standard error of the mean.

Results for the amplitude comparison of AB vs. CB comparison are presented in Figure 9, positive difference values indicate that amplitude was greater following tone A than tone C. During the ‘early’ window (10-40 ms post transition), tone B mean response amplitude (reported in mean amplitude in *pAm*^−1^ * 10^4^) is higher after tone A than C at the level of the left AC (mean: 9.55, SD: 21.43; Z = 2.06, *p* = .025); right AC (mean: 31.33, SD: 39.46; Z = 2.20, *p* = .00038); IC (mean: 7.05, SD: 11.48; Z = 2.96, *p* = .0038), and at the CN (mean: 5.22, SD: 10.89; Z = 2.63, *p* = .0070). It was not significant at the MGB (mean: 1.13, SD: 6.53; Z = 1.32, *p* = .24).

**Figure 9:**
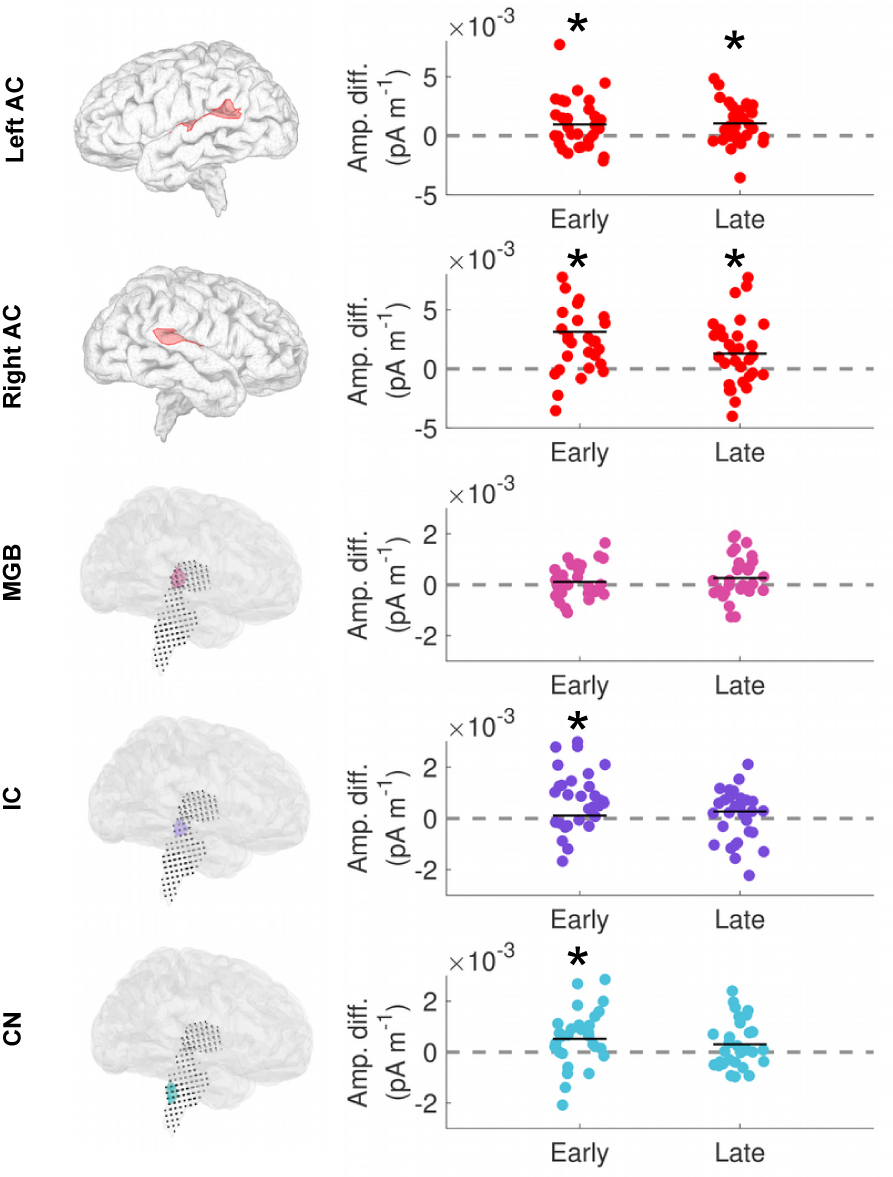
Effect of preceding stimulus on the amplitude of response to tone B, for each region of interest. An effect is found during the early window (10-40 ms) for most levels of the auditory system, but only the cortex shows a sustained effect into the late window (40-70 ms). Dashed grey lines indicate no difference between response amplitudes when tone B is preceded by A vs. C. Solid black lines show the group mean. Statistically significant comparisons are indicated with an asterisk (⋆), and are FDR corrected for multiple comparisons.

A difference between conditions was also observable within the later window (40 - 70 ms post transition), but only at cortical levels: left AC (mean: 12.85, SD: 28.22; Z = 2.88, *p* = .01); right AC (mean: 12.85, SD: 28.22; Z = 2.20, *p* = .035). Subcortical levels did not show significant differences: MGB (mean: 2.68, SD: 8.41; Z = 1.32, *p* = .12), IC (mean: 1.89, SD: 11.80; Z = 0.80, *p* = .21), and at the CN (mean: 3.06, SD: 9.25; Z = 1.42, *p* = .13). Reported P-values are FDR-adjusted within each analysis.

## 7. General Discussion

The main aim of this work was to establish evidence for true oscillatory entrainment in the auditory system. Consistent with this goal, Experiment 1 showed that MEG- and EEG-based FFR frequencies displayed an oscillatory aftereffect for several cycles post stimulus offset at each level of the auditory system (CN, IC, MGB, and left and right AC, based on MEG localization; Figures 1 and 2). Also consistent with the oscillation hypothesis, tracking of the FFR fundamental frequency showed that it converged to the stimulus frequency from a lower value over about 100 ms (corresponding to about 9 cycles of the F0), and then progressively diverged back to a lower value after stimulus offset (Figures 3 and 4; Table 1). Experiment 2 provided further evidence for oscillatory interactions between the offset of one stimulus and onset of the next. Tracking the FFR frequency of the lowest tone used, 98 Hz, showed that it was systematically higher for the first 30 - 40 ms when preceded by higher-frequency tones (Figure 7). Finally, transitions to the same tone from a higher- or lower-frequency preceding tone influenced the FFR amplitude of the target tone for the first 30 ms, such that amplitude was higher when the preceding FFR amplitude was also higher; this effect was seen at both cortical and subcortical levels, but appeared to persist longer at cortical levels (Figure 9).

Declaring the existence of neural entrainment phenomena must be approached with caution because many apparent oscillatory phenomena can be due to other factors (Haegens and Golumbic, 2018; Gourévitch et al., 2020; Obleser and Kayser, 2019). Haegens and Zion Golumbic propose three stringent criteria. The first criterion is the continuation of oscillatory activity for at least a number of cycles beyond external input (i.e. reverberation); results from experiment 1 clearly support the existence of a reverberation or aftereffect, and results from experiment 2 show that it interacts with incoming stimuli, leading to measurable differences in the frequency and amplitude of their representation. The search for ‘true’ oscillatory entrainment in the auditory system and their role has been confounded by methodological issues in the past (Gourévitch et al., 2020). In the present work, we studied a well-characterized auditory evoked brain response, and selected analyses like frequency tracking that are unlikely to introduce artifactual oscillations (as might narrow-band filtering or broad spectral analysis). Our data are consistent with prior reports of poststimulus FFR aftereffects in EEG data (Xu and Ye, 2015), and with recent data also documenting similar aftereffects from intracerebral electrode cortical recordings and cortical MEG in humans in the 60-80 Hz range, similar to the stimuli used here (Lerousseau et al., 2019; Ross et al., 2020). Taken together, these convergent findings provide strong evidence for entrainment, and our data furthermore indicate that it is a pervasive feature of auditory processing in subcortical nuclei as well as cortex. The lack of clear evidence for convergence in the EEG data points to the importance of separating the different sources of FFR throughout the auditory system, and the complementary nature of the MEG and EEG signals (Ross et al., 2020).

The second criterion for neural entrainment is phase alignment, which occurs when external information is introduced that contains frequencies close to the oscillator’s intrinsic rate, after which dampening or relaxing to the initial state should occur. Frequency-tracking results from both experiments support a gradual phase alignment or convergence of the brain’s frequency representation towards the stimulation frequency, and in experiment 1, post-offset relaxation. This finding fits with the concept that incoming stimuli engage an endogenous oscillatory mechanism, as has been suggested in lower frequency ranges (Thut et al., 2012).

The third criterion for entrainment is endogenous oscillatory activity in the absence of stimulation. Although we cannot observe endogenous activity in the absence of stimulation using these time-locked averaging-based techniques, the onset frequency may indicate sub-threshold non-aligned activity, at the system’s preferred frequency. In experiment 1, in which silence preceded the tone, tracking frequencies in the cortex started well below stimulation frequency, suggesting an oscillator with a lower preferred frequency. More work is needed to establish whether there are multiple preferred frequencies within the auditory system, or whether the stronger FFR responses observed at some frequencies are caused by constructive interference of the feed-forward signals (Tichko and Skoe, 2017).

An important open question is whether and how entrainment of oscillations plays a functional role in the auditory system (Haegens and Golumbic, 2018; Gourévitch et al., 2020; Thut et al., 2012). Entrainment in lower frequency ranges (i.e. delta, theta bands) has been proposed as a mechanism during active stimulus processing, for example to define the parsing window of linguistic segments from continuous speech (Kösem et al., 2018; Gourévitch et al., 2020). The FFR is an evoked response to a periodic signal, but it is not known whether there is some advantage to an oscillatory representation as opposed to a veridical feed-forward one. The FFR clearly carries periodicity information, and its strength and quality is associated with performance on perceptual tasks with a pitch component, including hearing-in-noise tasks (Li and Jeng, 2011; Presacco et al., 2019; Bharadwaj et al., 2015; Marmel et al., 2013; Anderson et al., 2010; Coffey et al., 2017; Song et al., 2011), suggesting that resistance to noise or sound degradation could be one of its functions. The oscillatory aspect might act as a very fine-scale temporal predictor, such that an incoming impulse of sound falling on the next cycle would contribute to its strength, and small timing differences could entrain the oscillatory as evidence of the external cycle length accumulates over cycles. The apparatus that produces periodic sounds (e.g. vibrating vocal chords, strings or columns of air) are constrained by physical properties and generally continue for multiple cycles, making frequency stability ecologically relevant.

The frequency tracking data indicate that the FFR is surprisingly sluggish, at least at the frequency tested, to reach the veridical value corresponding to the fundamental frequency of the stimulus. However, psychophysical studies show that pitch discrimination and identifications improve as a function of duration up to and even beyond a range of 100 ms, similar to the duration required for the FFR to reach the nominal frequency of the fundamental recorded in the present study (see Freyman and Nelson (1986); Robinson and Patterson (1995); Micheyl et al. (2012) for a review). Thus, it is reasonable to propose that the oscillatory mechanism is responsible for the fact that pitch processing improves with more cycles of input. There were large individual differences in the accuracy of the FFR tracking in our data, however, as well as differences between cortical and subcortical FFR sources; it will therefore be interesting to see how this variability is related to individual differences in pitch perception, which can also be quite large (Micheyl et al., 2006).

In contrast to models for syllabic processing, for example, in which a distinction is made between predictive coding (‘what’) and predictive timing (‘when’) (Arnal and Giraud, 2012), in the case of frequency coding, the distinction may not apply. We therefore speculatively propose a second function: that the oscillator’s ongoing activity might act as an internal template to which incoming information can be compared; in effect, the ‘what’ predicted by the system is pitch continuity. A low-level prediction mechanism that could rapidly generate error signals when stimulus periodicity changes could be useful, because encoding rapid changes in auditory information is critically important in complex sound processing, as in speech or music. Future work is needed to determine whether differences between incoming information and the FFR oscillatory entrainment produce brain responses consistent with predictive coding signals (Gourévitch et al., 2020; Chao et al., 2018).

If the FFR is generated by an oscillatory entrainment mechanism, as shown here (in both the MEG- and EEG-FFR; Figure 5), the future challenge will be to understand its relationship to behavioral variables. FFR amplitude and fidelity are known to correlate with many behavioral functions including fine pitch discrimination, hearing-in-noise perception, reading, and musicianship (reviewed in Coffey et al. (2019); Kraus et al. (2017)). Because the strength of the after-effect is strongly correlated with that of the FFR steady-state, and because the FFR includes an oscillatory component, observed behavioural correlations are suggestive of a functional role of entrainment, but cannot be conclusively separated from the quality of feed-forward encoding. Neither can the longevity of the aftereffect be necessarily used as a metric of strength of oscillatory entrainment, as protective mechanisms that prevent the development of hypersynchronicity may be independent of the oscillatory mechanism itself. For example, cross-frequency interactions with downstream regions may contribute to adjusting phase and frequency of local oscillators flexibly, depending on context and internal prediction models (Baillet, 2017) - a mechanism that appears to be impaired in epilepsy (Samiee et al., 2018). Clever approaches will be needed to investigate the perceptual function; for example, Baltus and Herrmann showed that the ability to perceive short gaps in tones is linked to an individual’s preferred auditory steady-state response (∼40 Hz; Baltus and Herrmann 2015), suggesting that an oscillatory response in lower frequency ranges might influence perception. Similar approaches could be used to clarify the perceptual relevance of individual differences in FFR entrainment mechanisms.

Our results establish that frequency coding in the human brain includes a mechanism of oscillatory entrainment, with the strongest effects at the cortical level, but found throughout the system. More work will be needed to establish how this mechanism arises as a cumulative product of circuits and even cells with resonant properties at or between different subcortical and cortical levels. This work may lead to a better understanding of neural signal processing principles underlying both low-level behavioral properties such as pitch discrimination, as well as higher-order cognitive mechanisms of speech and music perception.

## 8. Acknowledgements

This work was supported by a Foundation grant from the Canadian Institutes of Health Research to RJZ. We would like to thank undergraduate Annie Qin for assisting with piloting, and the team at the McConnell Brain Imaging Centre (Montreal Neurological Institute) for supporting and making accommodations for the study, particularly Elizabeth Bock; and the Brainstorm team.

## 9. Author contributions

E.B.J.C and R.J.Z. designed research; E.B.J.C., I.A.-B., and X.Z. collected data; E.B.J.C, I.A.-B. and X.Z. analyzed data with guidance from S.B.; and E.B.J.C. and R.J.Z. wrote the paper with contributions from all authors.

